# Interactions of medial and lateral prefrontal cortex in hierarchical predictive coding

**DOI:** 10.1101/439927

**Authors:** William H. Alexander, Thilo Womelsdorf

## Abstract

Cognitive control and decision-making relies on the interplay of medial and lateral prefrontal cortex (mPFC/LPFC), particularly for circumstances in which correct behavior requires integrating and selecting among multiple sources of interrelated information. While the interaction between mPFC and LPFC is generally acknowledged as a crucial circuit in adaptive behavior, the nature of this interaction remains open to debate, with various proposals suggesting complementary roles in (*i*) signaling the need for and implementing control, (*ii*) identifying and selecting appropriate behavioral policies from a candidate set, and (*iii*) constructing behavioral schemata for performance of structured tasks. Although these proposed roles capture salient aspects of conjoint mPFC/LPFC function, none are sufficiently well-specified to provide a detailed account of the continuous interaction of the two regions during ongoing behavior. A recent computational model of mPFC and LPFC, the Hierarchical Error Representation (HER) model, places the regions within the framework of hierarchical predictive coding, and suggests how they interact during behavioral periods preceding and following salient events. In this manuscript, we extend the HER model to incorporate real-time temporal dynamics and demonstrate how the extended model is able to capture single-unit neurophysiological, behavioral, and network effects previously reported in the literature. Our results add to the wide range of results that can be accounted for by the HER model, and provide further evidence for predictive coding as a unifying framework for understanding PFC function and organization.

## Introduction

Medial and lateral prefrontal cortex (mPFC/lPFC) are core hubs of the cognitive control and decisionmaking network in the brain (Cole & Schneider, 2007). The regions are densely and reciprocally connected (Barbas & Pandya, 1989; Barbas & Rempel-Clower, 1997), suggesting that their contribution to behavior depends in part on their tightly-coupled interactions during preparation, execution, and monitoring of the consequences of actions. Although these regions have long been the target of focused investigation, it remains an open question as to how they collaborate in an ongoing and interactive fashion to support adaptive behavior(Badre & Nee, 2018).

Independently, both regions have been implicated in a wide range of functions, and these functions appear to suggest a dissociation between the regions along a temporal dimension. Activity in mPFC, especially dorsal anterior cingulate cortex (dACC), is frequently observed in conjunction with behaviorally-salient events, such as error commission (Gehring, Coles, Meyer, & Donchin, 1990; Shen et al., 2015) or the appearance of stimuli cueing multiple, potentially conflicting, responses (Brown, 2009). In contrast, activity in lPFC is typically associated with long-term behavioral contingencies, such as maintaining information over extended periods of time(Sawaguchi & Goldman-Rakic, 1991) and representing the structure of ongoing tasks (Badre & D’Esposito, 2007). This temporal dissociation has been interpreted as reflecting complementary aspects of task performance in which mPFC signals changes in behavioral requirements, including specification of control signals (Shenhav, Botvinick, & Cohen, 2013) or selection of action policies(Holroyd & Yeung, 2012), and the implementation of appropriate control measures is delegated to lPFC (Botvinick, Braver, Barch, Carter, & Cohen, 2001).

How such temporally specific functional contributions of medial and lateral PFC can give rise to a wide variety of cognitive and behavioral effects has been formalized within the framework of hierarchical predictive coding(Alexander & Brown, 2015, 2018). The Hierarchical Error Representation (HER) model states that error signals generated by mPFC are used to train *error representations* in lPFC, and that, once learned, error representations maintained by lPFC serve to contextualize subsequent error calculations carried out by mPFC (Figure 1). Using this basic circuit as a repeating computational motif, the HER model is able to learn complex cognitive tasks in a manner that accords with human behavior, and measures of activity derived from error calculation and representation in the model reproduce qualitative patterns of single-unit and neuroimaging data.

**Figure 1.**
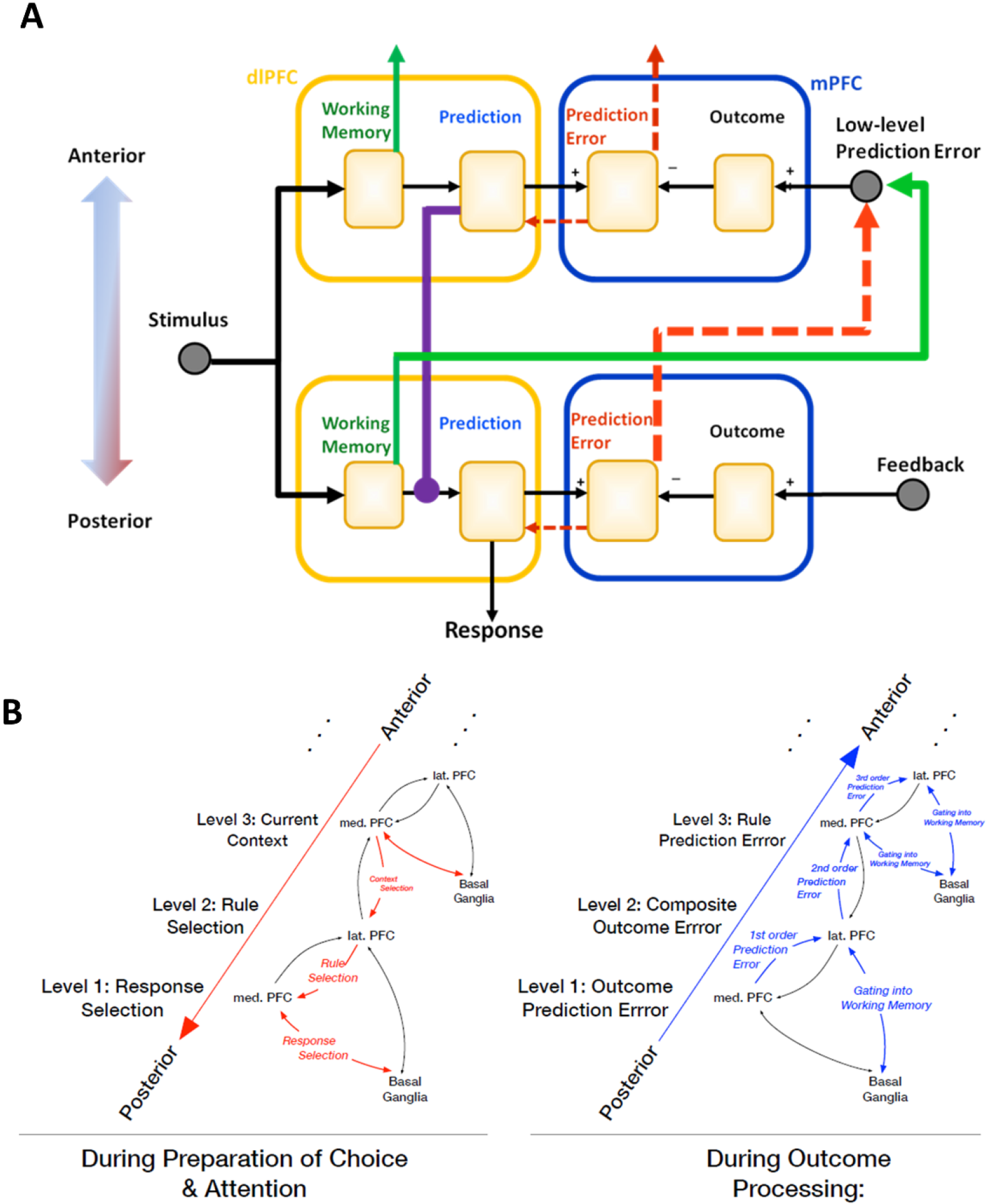
Interactions of mPFC and lPFC in the HER model. **A)** The HER model is organized hierarchically, with each hierarchical level instantiating a computational motif of prediction and prediction error calculations. Information flows between hierarchical levels along top-down and bottom-up pathways, which carry information regarding likely errors in a given context (top-down), and error signals (bottom-up) derived from external feedback (base hierarchical level) conjoined with items maintained in working memory (additional hierarchical levels). **B)** Reciprocal connections between mPFC and lPFC correspond to a putative hierarchical rostrocaudal gradient (Badre & D’Esposito, 2009) with bottom-up and top-down pathways governing the adjustment of behavior during ongoing task performance. **C)** During a trial, top-down connections (left) serve to establish the current context and relevant rules that govern eventual behavioral responses during preparatory periods, while following the generation of a response (right), bottom-up error signals are used to update outcome predictions and derive composite “proxy” outcomes that train higher-order representations of rules and contexts

While the HER model suggests how and when mPFC and lPFC might interact, our previous implementation of the model was aimed at the “event” level (Alexander & Brown, 2015): model equations were applied, and activity derived, following the occurrence of salient behavioral events, such as the presentation of task stimuli or performance feedback. The HER model is therefore only able to account for mPFC/lPFC interactions at a temporally coarse level, and is unable to capture the co-evolving development of activity during intra-event periods, nor can it capture aspects of behavior, such as switch costs (Wylie & Allport, 2000), that may manifest through differences in response time. Thus, although the HER model provides a promising framework for understanding the function, organization and interactions of PFC regions (Alexander & Brown, 2018), a significant gap in its explanatory power remains. In this manuscript, we present an extended version of the HER model that incorporates real-time dynamics. Simulations of the extended model show how reciprocal activity derived from the model can account for additional effects observed from behavioral, single-unit, and brain network data, and consequently provides further support for a predictive-coding interpretation of PFC function.

### Methods

The operation of the HER model is described in detail in previous publications (Alexander & Brown, 2015). Generally, the HER model is composed of 2 or more hierarchical levels, and each level of the hierarchy is made up of three primary components: a working memory (WM) store **r**, and weight matrix **X** determining the probability that a stimulus **s** will be stored in **r**, and a weight matrix **W** determining how items stored in WM either influence behavior (at the lowest hierarchical level) or modulate the processing of lower hierarchical levels. Gating into working memory is determined by the pair of equations:

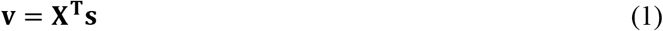

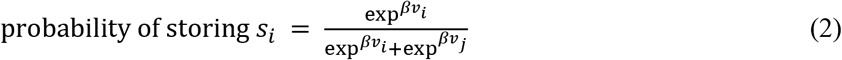

Eq. 1 reflects the learned association between task-related stimuli and prediction errors generated at each hierarchical level of the model. Eq. 2 is a softmax function calculating the probability of storing the current stimulus s_i_ in WM relative to the currently contents of WM, indexed by *j*, and β is a gain parameter governing the degree by which differences in elements of **v** influence the probability of maintaining or updating WM. Intuitively, these equations determine which of multiple stimuli is more “valuable” to store in WM (eq1), and to transform this value into a probability (eq2). Once stored in WM, the representation of a stimulus is used to generate the output of that hierarchical level:

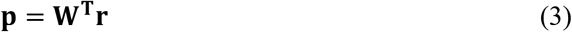

At the lowest hierarchical level, network output is used to generate a behavioral response using another softmax function:

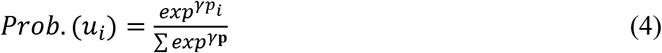

where γ is a gain parameter indicating how likely the model is to select the response with the highest value. The output of higher-order hierarchical levels, **p’**, is used to modulate the processing of lower-order levels output as follows:

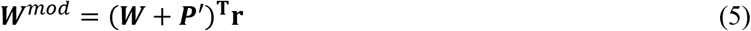

Here, **P’** is the output **p’** of the higher-order hierarchical level reshaped in a matrix with the same dimensionality as **W**.

#### Learning

Learning in the model depends on the calculation of error signals at each level of hierarchy. At the base level, errors are computed as:

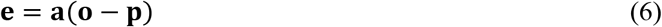

where **o** is a feedback vector indicating the response identity and whether the response was correct or incorrect, and **a** is a filter that is 1 for the index of the generated response and 0 everywhere else. Associative weights at each level are updated according to:

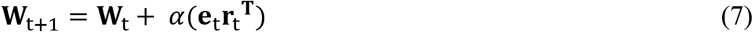

and α is the learning rate. To train higher-order hierarchical levels, a proxy outcome signal is composited from the error signal computed at the lower level (Eq. 6) and the active WM representation at the lower level. This is done by taking the outer product of the lower-level error and representation vectors:

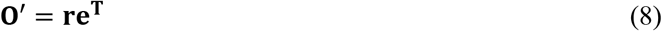

For computational convenience, **O’** is reshaped into a vector **o’** with the same dimensionality as **p’.** Training of WM weights **X** is done by backpropagating the error term **e** for each level:

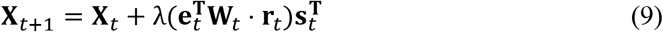

with learning rate λ.

#### Simulated task

The HER model was developed to account for behavior and brain function in tasks requiring the selection and application of rules that govern how to respond to a concrete stimulus (e.g., an arrow cueing response identity). We therefore selected a recent, rule-based task(Oemisch et al., 2018) in which, on each trial, monkeys were presented two stimuli, one on each side of the screen (Fig. 2A). Stimuli had two behaviorally-relevant features that varied independently from each other, namely color (red/green), and the motion direction of a visual pattern (up/down). The task requires that the monkey learn which of the two colors is currently relevant, and to respond with a saccade in the up- or downward motion direction of the stimulus with the currently relevant color. Once the relevant color has been learned, successful performance of the task requires that, on each trial, the monkey first identifies the location (side of the screen) of the appropriately-colored stimulus, after which the identity of the response (up/down) corresponding to the direction of the pattern movement is determined. The task described in Oemisch et al. additionally incorporates periodic reversals when the relevant color changes and the monkey must adapt their responses to the new task contingencies; while the HER model is able to learn such reversals (cf. Fig. 2C), this aspect of the task is not further considered here as it does not bear directly on the *within-trial* temporal dynamics of PFC interactions.

**Figure 2.**
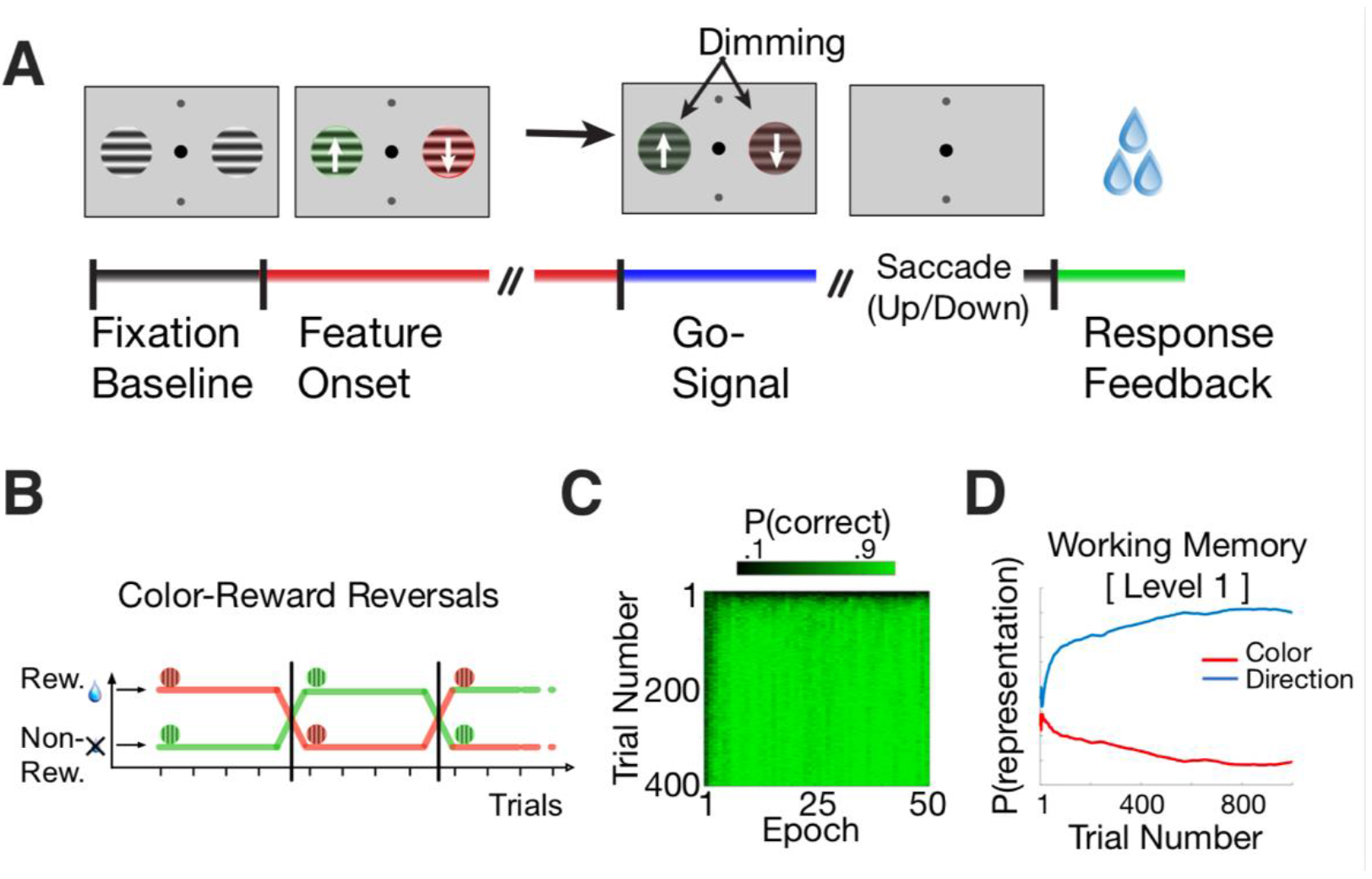
**A)** The color-based reversal task described in Oemisch et al. (2018). Subjects are shown a fixation point and two neutral stimuli. Then the stimuli switched to oppostie colors and began to move within their apertures in opposite, upward or downward directions of motion. Following a brief interval, a ‘Go’ (dimming) signal is presented, following which the subject is required to indicate the direction of the grid for the currently relevant color. The dimming signals occurred at unpredictable moments in time either in both stimuli simultaneously or in sequence to control covert attention (not shown). **B)** The rewarded, relevant color reversed uncued after ≥30 trials, and subjects had to adjust their behavior to indicate the movement direction associated with the newly relevant color. **C)** The HER model is able to learn the Oemisch et al. reversal task easily. During an initial period lasting from 1-10 reversal epochs, the model learns to preferentially gate in task features to hierarchical levels. Following reversals, the model rapidly learns the new task contingencies while preserving the hierarchical order of information. **D)** The concrete decision variable in the reversal task is the apparent direction of motion associated with the currently relevant color. The HER model learns to represent this variable at the lowest hierarchical level (Level 1), consistent with its direct relevance to generating behavioral responses.

#### Model Simulations

To simulate real-time dynamics in the HER model, we adopt the approach taken by previous studies of network models of cognitive control (e.g., (Yeung, Botvinick, & Cohen, 2004)) in which the process used to establish weights suitable for performing a task is carried out independently of simulations of temporal dynamics. In order to establish model weights suitable for performing the Oemisch et al. reversal task, a previously-described version of the (event-level) HER model (Alexander & Brown, 2015) using the same parameters was trained on 20,000 trials of the reversal task, divided into 50 blocks of 400 trials each. On each trial during learning, 3 events were modeled: the onset of task stimuli, the occurrence of a response and feedback, and a neutral cue indicating the start of the intertrial interval (ITI). Stimuli were modeled as a binary vector, with elements corresponding to color and movement direction (four total elements per stimulus), and independent stimulus vectors were used to model each side of the simulated display, for a total of 8 task stimulus elements. A 9^th^ element was used to indicate the ITI. The model was permitted 3 responses, two corresponding to movement direction (up/down), and one neutral response indicating acknowledgement of ITI onset. Each response could be associated with two outcomes (correct/incorrect), for a total of 6 response-outcome predictions. Responses were generated by subtracting, for each response, the prediction of incorrect feedback from the prediction of correct feedback, and passing the values through a softmax function, as described in the methods section.

#### Temporal Dynamics

The network weights recovered from the training procedure were fixed during the real-time simulations as described above. In order to simulate real-time dynamics in the network, changes in unit activity on each cycle were modeled by a non-linear “shunting” equation (Grossberg, 1988):

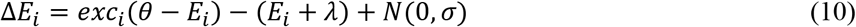

where θ is the upper asymptotic activity a unit could achieve (set to a value of 10), λ is the lower boundary toward which unit activity decays passively (−0.05), and N is gaussian noise applied to the signal change with mean 0 and variance σ = 0.01. *E*_*i*_ is the current activity of unit i, and *exc*_*i*_ is the current net excitatory input to the unit, computed as in equations 1,3 & 6. Specifically, exc_i_ is equal to **v** in eq. 1, governing the dynamics of WM update and maintenance. For computing model predictions at all hierarchical levels, exc_i_ is equal to **p** in eq. 3. Finally, for computing error (eq. 6), exc_i_ is set to **e**, the difference between predictions **p** and observed outcomes **o**.

Each simulated trial lasted up to 700 cycles, and on each cycle the processing steps outlined in eqs. (1-9) were followed and unit activity updated according to eq. 10. During the first 100 cycles of the trial, no input was presented to the network. On cycle 101, a stimulus vector representing the color and movement direction of the left and right stimuli was presented to the network, after which the network was able to register a response. Following a response, the stimulus vector was set to 0 and the network was provided feedback for 50 cycles. Following the offset of feedback, the network was run for an additional 250 cycles prior to the beginning of the next trial. Network activity was not re-initialized after each trial. As described in previous work(Alexander & Brown, 2011, 2014, 2015, 2018), eqs. 1–9 specify two primary signal types, namely prediction (eqs. 1 & 3) and error (eq. 6). In the simulations reported here, both types of signals are subject to the temporal dynamics embodied in eq. 10.

## Results

The HER model was able to learn and perform the reversal task relatively easily. During an initial “burn-in” period (Fig. 2C), the model primarily learned to hierarchically segregate relevant feature dimensions (Fig. 2D): the WM gating mechanism (eqs 1&2) learned to store the “concrete” movement direction at the lowest hierarchical level while color information was stored in the superior hierarchical level. This pattern accords with intuitions regarding how individuals might solve the task (Oemisch et al., 2018): color acts as a ‘rule’ that indicates which concrete stimulus (movement direction) should govern the ultimate response. After this initial learning period, the model is able to perform reversals within a short period of time. Because the model has learned stable WM mappings, learning to respond to the opposite color entails relatively rapid changes in top-down modulation of concrete responses, i.e., rather than relearning the task from the ground up. These findings suggest one manner in which hierarchical representation of information might support rapid and flexible reconfiguration of responses in the face of changing task contingencies.

### Response Preparation

Introducing real-time dynamics allows us to investigate the evolution of predictive activity in the model during a trial, the influence of previous trial effects, and derive measures of reaction time from model activity (Fig. 3). During preparatory periods following the onset of a task stimulus the activity of predictive units in both mPFC (Fig. 3A) and lPFC (Fig. 3B) begins to ramp up. Normalized predictive activity over the first 200 cycles of a trial (Fig 3C) suggests a causal relationship of lPFC to mPFC: activity in lPFC increases more quickly early in the trial, particularly when the target response direction changes. The development of activity depending on previous trials additionally influences the speed at which a response is generated (Fig. 3D) due to lingering model activity related to features from the previous trial. When both the position of the relevant stimulus and the cued response remain the same, model reaction times are most rapid; activity in both lPFC and mPFC starts above a resting baseline, making it easier for response signals to exceed a threshold (eq. 10). In the model lPFC, this starting point is primarily sensitive to changes in the location of the relevant stimulus; when the location changes, lPFC activity representing the prior location must first be suppressed prior to the representation of the new location becoming active and able to modulate mPFC activity. In contrast, mPFC activity is sensitive to changes both in the location of the target stimulus, as well as changes in the cued direction; both factors influence the development of mPFC activity, but here changes in the cued response tend to have a greater influence – as noted above, the cued response direction is a concrete decision variable that directly drives model actions, while identifying the location of the relevant color stimulus acts as a rule that modulates ongoing response activity. Artificially lesioning lPFC (Fig. 3E) prevents this modulation, after which the development of mPFC activity is only influenced by differences in the cued response direction. RTs derived from the model (Fig. 3D) replicate standard trial sequence effects wherein feature repetition facilitates responding, while feature switches interfere with responses(Fecteau & Munoz, 2003). The HER model further suggests why some feature switches may produce greater interference effects than others – specifically, switches of the feature that most directly drives response results in a greater RT difference than changes in the more abstract feature dimension.

**Figure 3.**
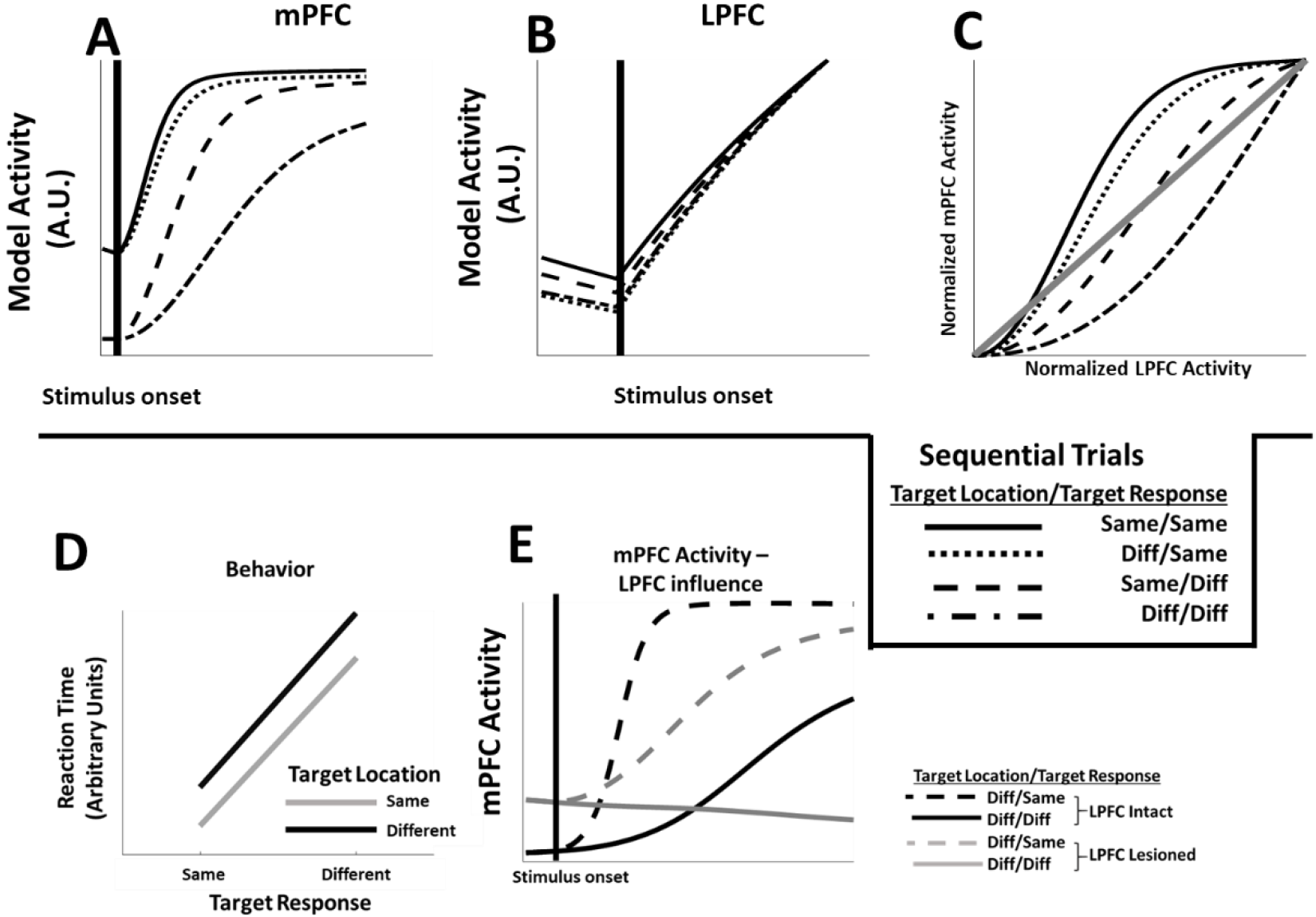
**A)** MPFC activity in model simulations is influenced by switches or repetitions of feature dimensions. When both feature dimensions (location & direction) repeat, as well as repetitions of the target direction, mPFC activity reaches asymptote quickly after stimulus onset, while changes in the target response produce delays in the evolution of mPFC activity. **B)** Activity in LPFC likewise shows intertrial effects; however, in this case, delay in the development of LPFC activity is due principally to shifts in the location of the target stimulus, while changes in the more concrete target direction variable have relatively little influence on LPFC activity. **C)** LPFC activity develops more quickly than mPFC activity during early trial stages, and continues to increase after mPFC activity asymptotes, consistent with a role for lPFC in modulating mPFC responses. **D)** The delay in model activity following feature switches contributes to changes in model reaction times: reaction times are most rapid on trials in which both feature dimensions repeat, while switches in either or both features result in longer reaction times. **E)** Artificially lesioning the simulated LPFC component of the model further delays the development of mPFC activity in the model, demonstrating a causal influence of LPFC on mPFC function in the HER model.

### Feedback Processing

Following a response, the model receives feedback indicating whether the selected response was correct or incorrect. While equations used to compute errors in the model, like those used to calculate prediction unit activity, apply at every moment in the simulations, error-related activity is most prominent following feedback delivery, during which ongoing predictive activity is compared to an experienced outcome. During learning in the event-level model, comparison of feedback and concrete outcomes occurs only at the lowest hierarchical level; at superior hierarchical levels, “outcomes” are derived from WM representations at the inferior level combined with the results of the feedback comparison process (i.e., the error signal). These “proxy” outcomes constitute a higher-order training signal that is composited from lower-level WM representations and error signals; the composition of the higher-order training signal is carried out by lPFC in the HER model, while the comparison of outcomes and predictions is undertaken by mPFC.

Naturally, since the proxy outcome depends on the lower-level error signal, the evolution of error units in mPFC in our simulations precedes the development of activity in lPFC training units (Fig 4A, top frame): the error signal ramps up rapidly at the onset of feedback, and decays quickly following feedback offset. In contrast, the lPFC training signal lags the mPFC error signal, and its activity is temporally blurred. Although the relative onset of the error and training signals is prefigured by the architecture of the HER model, the relative distribution (Fig. 4A, bottom panel) of the signals emerges only due to the temporal dynamics introduced in these simulations. This emergent pattern matches data recorded from monkey dACC and lPFC during performance of the reversal task (Fig 4B; Oemisch et al., 2018), and adds to the already considerable array of effects the HER model has been applied to (Alexander & Brown, 2015, 2018).

**Figure 4.**
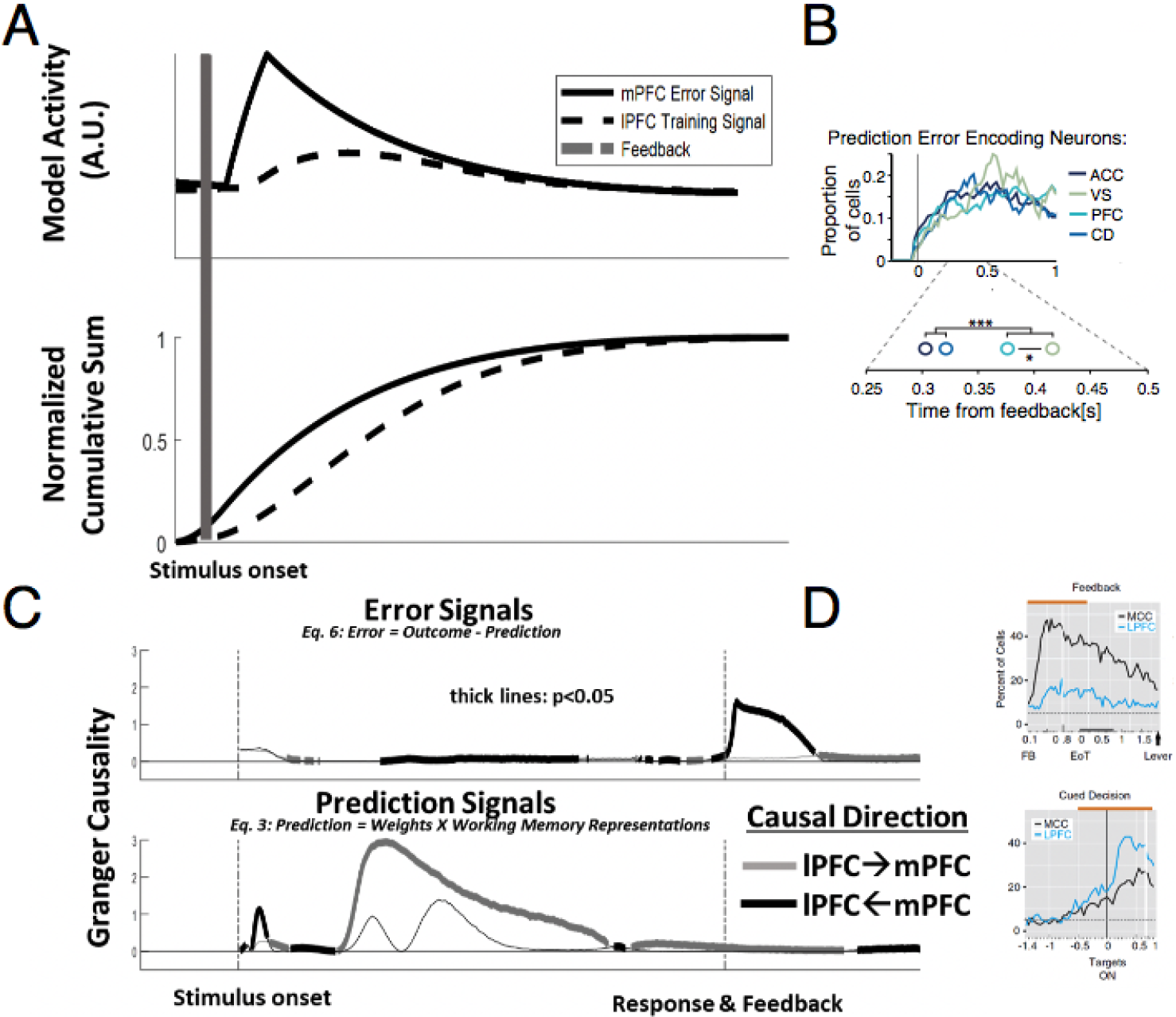
Temporal dynamics of error and prediction. **A)** Performance-related error signals in the model mPFC at the lowest hierarchical level evolve rapidly following the onset of feedback. Proxy outcomes used to train superior hierarchical levels are a composite of error signals calculated by inferior levels and the contents of WM, with a consequent lag in the temporal profile and lingering activity. **B)** The relative temporal profiles of error and composite training signals matches those observed in monkey mPFC and LPFC, respectively, performing the reversal task. **C)** Analysis of causality in the model shows a transient causal relationship of mPFC to lPFC following behaviorally salient events such as feedback (top frame) or stimulus onset (bottom frame). This causal relationship is reversed during trial periods involving the preparation and execution of response (bottom frame). The dynamic shift in causal direction over the course of a trial matches similar patterns observed in monkey mPFC and lPFC during feedback processing and cued performance (right frames, Stoll et al., 2016).

### Information Flow

While our previous results are suggestive of how mPFC and lPFC might interact during preparatory and feedback periods of a trial, a key strength of introducing real-time dynamics in the HER framework lies in the capacity to probe how mPFC/lPFC interactions develop continuously over the course of a trial. To do so, we turn to Granger causality(Luo et al., 2013) as a measure of how well one variable can be predicted by lagged values of another variable – here, the variables are the unit activity in mPFC and lPFC and Granger causality indexes whether unit activity in one area is better predicted by the preceding unit activity from the other area than by its own past. Granger causality was computed for a time lag of 1 model cycle for both error-related and predictive units in the model, and trial timing was standardized – even if a response was indicated prior, the model was simulated for 300 cycles following the onset of a stimulus. Consistent with our discussion above, causally significant error signals were observed primarily following performance-related feedback (Fig 4B). Immediately following the onset of feedback, Granger causality for the influence of mPFC on lPFC was significant at a level of p<0.05, and remained so until feedback-related activity naturally decayed. LPFC also causally influenced mPFC at a significance level of p<0.05, but only after an extended delay following feedback.

Also consistent with mPFC’s role in immediate processing of salient stimuli, prediction signals in the model immediately following stimulus onset Granger-cause activity in lPFC (p<0.05). While the HER model contains no mechanism by which predictions in mPFC influence processing activity in lPFC, Granger causality only indicates whether one signal can be predicted by previous values of another signal. In this case, predictive signals in mPFC and lPFC are correlated, but mPFC signals develop more rapidly than lPFC signals, producing a significant Granger causality effect. Following this transient effect, Granger causality for the influence of lPFC on mPFC becomes significant during the delay period prior to the generation of a response, consistent with the role of lPFC in maintaining information and implementing control demands.

## Discussion

In this manuscript, we have described additional simulations of the HER model in which real-time temporal dynamics were introduced to the model. The results of these simulations provide additional perspective on how the activity of mPFC and lPFC, as components of a hierarchical predictive coding framework (Alexander & Brown, 2018), might develop and interact following salient task events, and how the relative direction of this interaction evolves over the course of a trial. Beyond simply exploring the dynamics of model activity, our simulations demonstrate how the HER model can further account for additional single-unit (Oemisch et al., 2018), behavioral (Wylie & Allport, 2000), and network effects (Stoll, Fontanier, & Procyk, 2016) previously reported in the literature.

At the level of neurons, the activity of single units in the HER model corresponding both to lPFC and mPFC, is observed to ramp-up following the onset of a task-relevant stimulus. Previous real-time models of mPFC (Alexander & Brown, 2011, 2014) have likewise shown units with ramping activity profiles, similar to those of reward-and error-predicting neurons in monkey mPFC (Amador, Schlag-Rey, & Schlag, 2000; Amiez, Joseph, & Procyk, 2006). The HER model, conceived as a temporally-coarse hierarchical extension to the PRO model (Alexander & Brown, 2015), was unable to replicate this pattern; by re-introducing real-time dynamics, and consequently recovering effects from the PRO model that depended on temporal processes, our simulations underscore that the principal role and computational mechanisms attributed to mPFC by the PRO model remain intact in the HER model. Furthermore, the simulations in this manuscript extend real-time processing to the registration of feedback and the development of error signals. The original PRO model (Alexander & Brown, 2011) was developed using temporal difference (TD) learning formulations(Sutton, 1988) in which the temporal profile of feedback signals (and subsequent error signals) was specified by the modeler (either as a punctate event or a box car profile). Here, the duration and magnitude of feedback signals is still modeler-defined, but the development of activity registering these signals is described by the same timing equations used to model all other unit activity in the model. By extending temporal dynamics to feedback, the simulations here are able to capture the temporal profile of error-signaling units recorded from monkey mPFC and lPFC, as well as the relative onset and decay of these signals (Fig 4).

Specifically, simulated error-related signals in mPFC are observed to peak and decay more rapidly than signals observed in lPFC, consistent with recent reports (Oemisch et al., 2018; Shen et al., 2015). The HER model explains this through the role of mPFC in training error representations in lPFC: error signals generated directly by feedback in the model are combined with active representations of task-stimuli to derive higher-order outcome and error signals, represented in lPFC. As the development of these higher-order signals is mechanically subsequent to direct error signals (Fig 1), the dynamics specified in eq. 10 dictate a later peak and lingering activity. Furthermore, the “proxy” outcome signals derived in lPFC are required for subsequent higher-order error calculations, suggested by the HER model to be carried out in hierarchically-superior regions of mPFC (Fig 4), and these error signals are subject to additional lag as lower-order error and training signal computations that support their calculation develop. The HER model thus provides a mechanistic explanation for the relative time course of error signals in caudal-rostral regions of mPFC (Polli et al., 2005).

Although computation of error and prediction-related signals is ongoing throughout our simulations, causality analysis over the entire course of a trial reveals that the net direction of information flow depends both on the type of information (prediction or error) computed, as well as the period within a trial. Following salient task events, such as stimulus onset or delivery of feedback, information in the model flows primarily from mPFC to lPFC, while during periods in which a salient event is expected but has yet to occur, information flows principally from lPFC to mPFC. This pattern maps well both to functional roles attributed to these regions, as well as the observed time course of interactions. Functionally, mPFC has long been associated with processing novel or behaviorally-relevant events, especially the occurrence of errors (Gehring et al., 1990) or otherwise surprising (Ide, Shenoy, Yu, & Li, 2013; Jessup, Busemeyer, & Brown, 2010) stimuli, while lPFC is implicated in slower processes involving information maintenance (Sawaguchi & Goldman-Rakic, 1991), representing task structure (Badre & D’Esposito, 2007), or implementing control in preparation for upcoming demands (Botvinick et al., 2001). This temporal dissociation, implied in the architecture of the HER model (Alexander & Brown, 2015) is made explicit in this manuscript, and the relative timing and flow of information in the model is consistent with human and monkey studies of PFC (Oemisch et al., 2018; Shen et al., 2015; Stoll et al., 2016; Taren, Venkatraman, & Huettel, 2011).

Finally, by introducing temporal dynamics, we were able to use the HER model to replicate sequential trial effects that are a staple of the cognitive control literature (Fecteau & Munoz, 2003; Wylie & Allport, 2000). Unsurprisingly, reaction times for the model are the most rapid for trials in which all features of the chosen stimulus are identical, and slowest for trials in which all features have changed relative to the previous trial. Of interest, however, are trials in which only one relevant feature (out of two in the current study) changes. In these cases, the identity of the changing stimulus can have differential effects on reaction time. The HER model solves structured tasks by decomposing stimulus dimensions hierarchically (Alexander & Brown, 2015): dimensions that are “concrete”, i.e., those that most directly inform the eventual response, are preferentially encoded at the lowest hierarchical level, while more abstract features are encoded at superior hierarchical levels. The simulations reported here suggest that changes in the concrete decision variable (in this case, the direction of the target response) may have a more profound influence on reaction time than changes in the more abstract variable. Recent work (Vassena, Deraeve, & Alexander, submitted) has begun to explore how interfering with the structure of a task through manipulations of presentation order might influence decision making and performance. The results of the present study suggest a complementary approach in which the differences in performance elicited through feature changes might be used to infer the representation of task structure.

In summary, the simulations of the extended HER model reported in this manuscript demonstrate that including temporal dynamics endows the model with additional explanatory power, and provides the basis for additional work investigating the function and interaction of regions within PFC, as well as how they contribute to behavior. More generally, the HER model, as an instance of predictive coding, suggests how additional regions in PFC may be organized(Alexander, Vassena, Deraeve, & Langford, 2017); specifically, the HER model is primarily concerned with how information is integrated in order to generate responses, but may have little to say about how information is acquired to begin with. It is possible that additional regions implicated in cognitive control may be integrated with the HER framework to describe information is actively selected and interpreted to assist in adaptive behavior.

